# Historian: accurate reconstruction of ancestral sequences and evolutionary rates

**DOI:** 10.1101/093161

**Authors:** Ian Holmes

## Abstract

Reconstruction of ancestral sequence histories, and estimation of parameters like indel rates, are improved by using explicit evolutionary models and summing over uncertain alignments. The previous best tool for this purpose (according to simulation benchmarks) was ProtPal, but this tool was too slow for practical use. Historian combines an efficient reimplementation of the ProtPal algorithm with performance-improving heuristics from other alignment tools. Simulation results on fidelity of rate estimation via ancestral reconstruction, along with evaluations on the structurally-informed alignment dataset BAliBase 3.0, recommend Historian over other alignment tools for evolutionary applications. Historian is available at https://github.com/ihh/indelhistorian under the Creative Commons Attribution 3.0 US license. Contact: Ian Holmes.

## 1 Introduction

Multiple alignments are used for several purposes in bioinformatics, only one of which is homology-directed protein structure prediction, yet this is exactly the application that has tended to dominate alignment benchmarks. For evolutionary applications, such as reconstructing trees or ancestral sequences, there is evidence that different alignment tools (along with different tool-assessment metrics) might be preferable (Löytynoja and Goldman, 2008; Westesson *et al.,* 2012a). Aligners that are optimized for detecting structural homology may not do so well at recovering information about substitution rates, whereas explicit statistical models of the sequence evolution process may do a better job at estimating these parameters. Furthermore, the ideal way to estimate evolutionary parameters is not to use a single point estimate of the alignment, but to sum over alignments as a “nuisance variable”. Almost no tools do this, with the exception of MCMC samplers (Westesson *et al.,* 2012b; Redelings, 2014).

Empirical studies suggest that, for the purposes of estimating molecular evolutionary parameters—such as indel rates (Westesson *et al.,* 2012a), dN/dS ratios (Redelings, 2014), or trees (Löytynoja and Goldman, 2008)— it is advantageous to use a statistical model of evolution and to treat alignment rigorously as a “missing data” problem. In a previous study, we sought to quantify systematic biases introduced into the estimation of indel rates, using a simulation benchmark (Westesson *et al.,* 2012a). The ProtPal program (Westesson *et al.,* 2012a), which models indel evolution using transducers—finite-state machines which can be multiplied together like substitution matrices (Bouchard-Côté, 2013)—introduced the least biases of the tools evaluated. Unfortunately, the implementation of ProtPal published with that benchmark was too slow for practical use.

Here, we present a clean reimplementation of the algorithm underlying ProtPal in a new tool, Historian, that can also estimate rates by summing over alignments. We report an assessment of the alignment accuracy on structurally-derived benchmarks, together with a simulation-based experiment to quantify the accuracy of indel rate estimation.

## 2 Methods

Historian combines established algorithms from several sources. Like ProtPal, Historian progressively climbs a tree from tips to root, building an ancestral sequence profile that includes suboptimal alignments (Lee *et al.,* 2002; Westesson *et al.,* 2012a) using a time-dependent evolutionary model (Rivas and Eddy, 2015). The guide tree is found by neighborjoining on a guide alignment constructed from the all-vs-all pairwise alignment graph, or from a sparse random connected subgraph (Bradley *et al.,* 2009). The guide alignment, which can also be supplied by another tool, can optionally constrain the progressive reconstruction, which is followed by iterative refinement.

Historian also implements the phylogenetic EM algorithm for continuous-time Markov chains (Holmes and Rubin, 2002), so substitution and indel rates can be estimated directly from sequence data. Since the method builds an HMM that captures suboptimal as well as optimal alignments— rather like a partial order alignment (Lee *et al*., 2002)—the program can also estimate rates in an “alignment-free” way (i.e. summing over alignments) simply by running the Forward-Backward algorithm on this HMM, and using the posterior counts to weight the phylo-EM updates.

The principal parameter determining Historian’s execution speed is the number of suboptimal alignments it retains while building the progressive ancestral reconstruction. Historian provides a command-line option -fast which substantially reduces this number of alignments. We here report results for both default- and fast-mode operation.

## 3 Results and Discussion

We first performed a simulation benchmark to assess Historian’s ability to reconstruct evolutionary parameters, compared to other tools. We based the simulation parameters on the evolutionary profile of the HIV/SIV GP120 envelope domain, as follows. We started with a tree made from an alignment of ten HIV and SIV GP120 domains (Figure 1). We used the tree to estimate the indel rates for the alignment, and simulated 100 different alignments on this tree using a sequence length of 500 amino acids (roughly the same as the GP120 domain) with the expected substitution rate (*σ*) set to one substitution per unit of time and the simulated insertion and deletion rates (*λ, μ*) set to the midpoint of the insertion and deletion rates estimated for GP120 (0.028 indels per unit of time). For a substitution model, we used the Dayhoff PAM matrix, which we judged not to give an unfair advantage either to Historian (which by default uses a matrix estimated from PFAM) or to Prank (which uses the WAG matrix).

**Figure 1:**
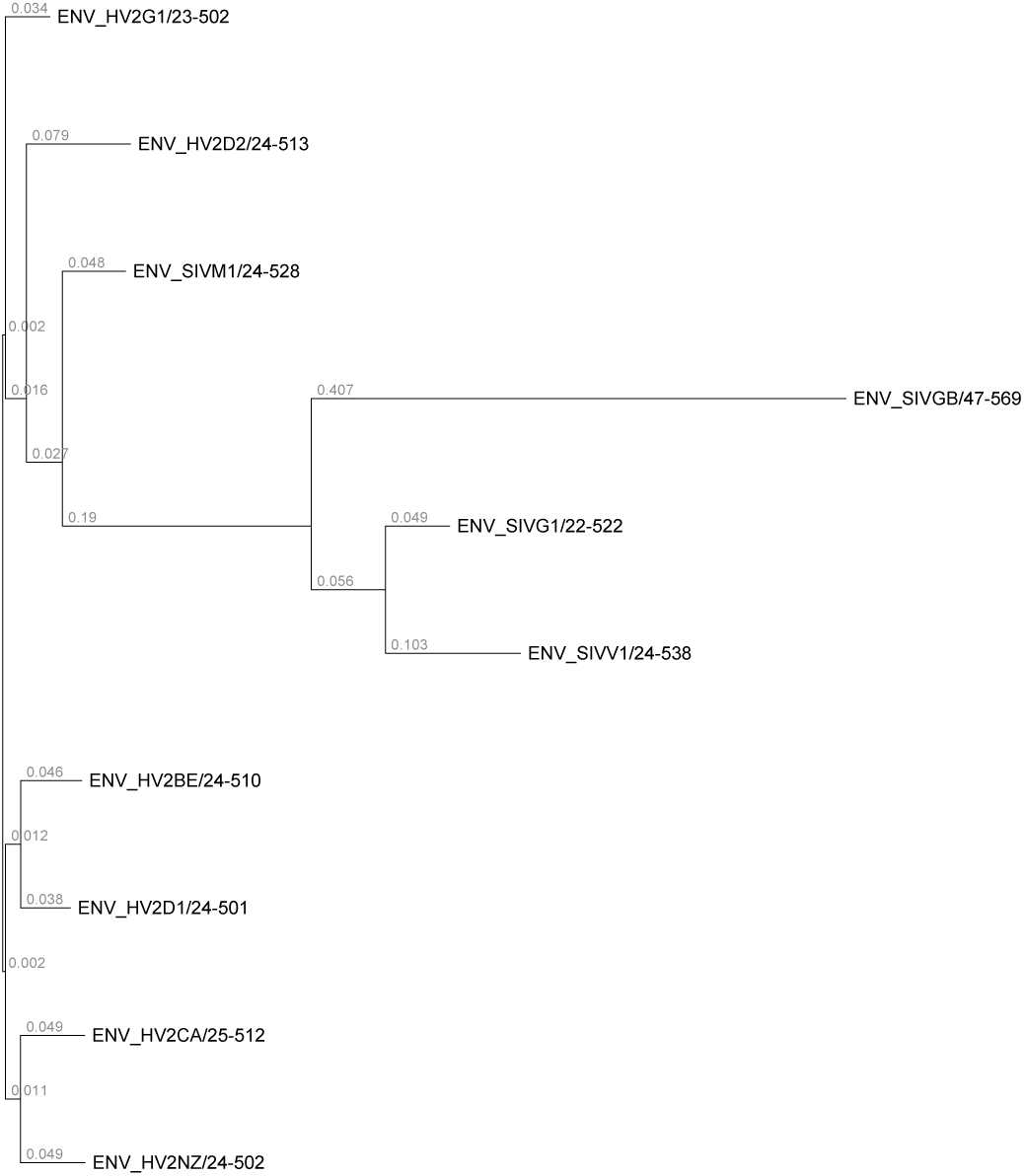
Tree of selected HIV and SIV GP120 domains, used for the simulation benchmark.

From the simulated sequence data, we estimated indel rates (a) using the true evolutionary alignment (supplied to Historian); (b,c) using Historian (alignment-free, default and fast modes); and (d,e,f) using Prank, ProbCons and Muscle, the best-performing tools from the ProtPal benchmark (Westesson *et al.,* 2012a). In each case, we estimated insertion and deletion rates 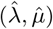 and computed relative errors 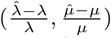 compared to the simulated indel rates. Having performed this experiment using the rates estimated for GP120 (*σ* =1, λ = *μ* = 0.028), we repeated the simulation, varying the rate parameters (*σ* = 2, λ = *μ* = 0.056; *σ* = 5, λ = *μ* = 0.14; *σ* = 2, μ = *μ* = 0.112; *σ* = 5, λ = *μ* = 0.28; *σ* = 5, λ = *μ* = 0.005; *σ* = 5, λ = *μ* = 0.01) while still using the tree of Figure 2. Varying rate parameters did not appear significantly to affect the ranking of the programs.

**Figure 2:**
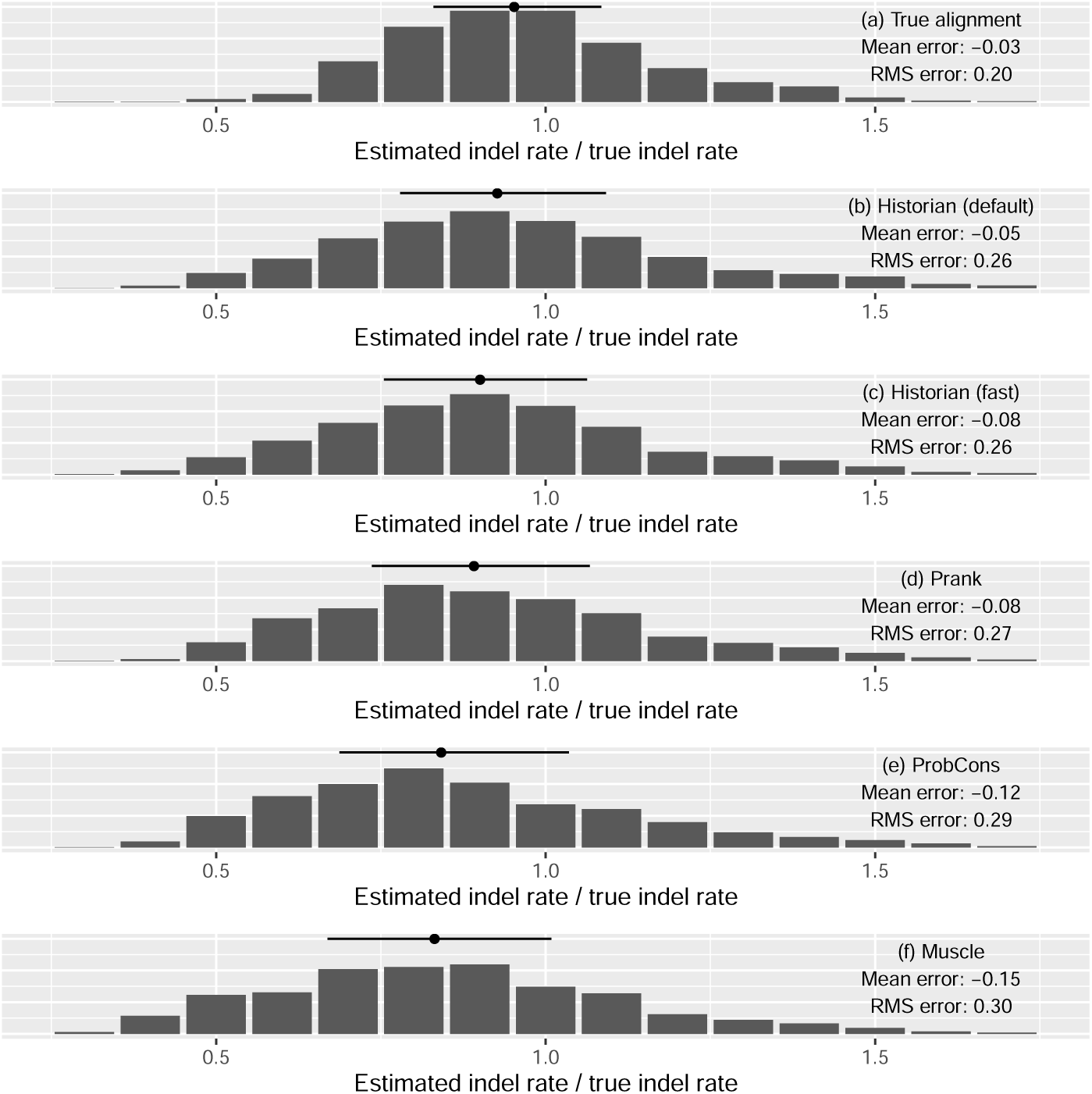
Results of the simulation benchmark. For each method evaluated, the distribution of the ratio of inferred to true indel rate is shown; for a perfect rate inference, this ratio would be equal to 1. The mean of the ratio is also shown; this represents the the systematic error, so e.g. using Muscle results in a systematic 15% underestimate of indels in these simulations. The root-mean-squared value of the ratio is also shown, and the median and interquartile range are annotated above each plot. Tool versions are as shown in Table 1.

**Table 1:**
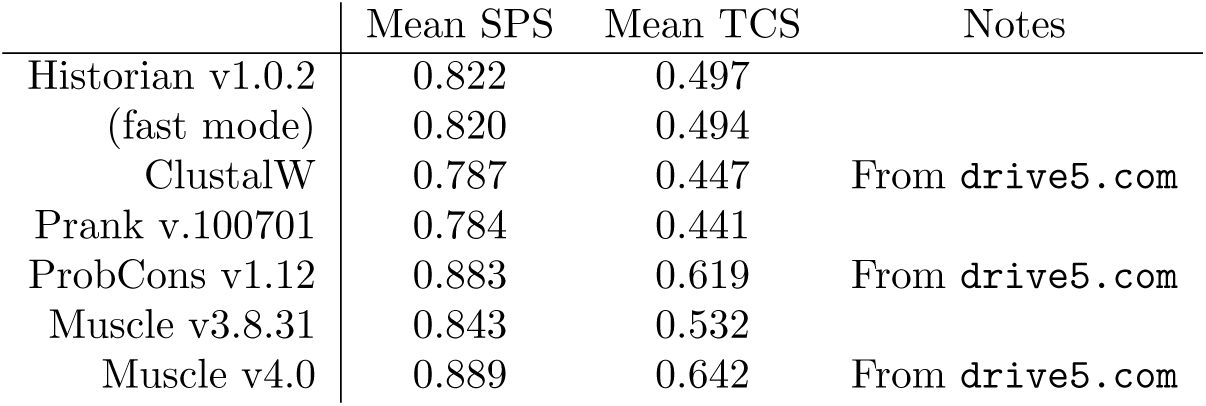
Comparison of Historian to other alignment programs using the BAliBase 3.0 benchmark with SPS and TCS alignment quality scores (Thompson *et al*., 2005).

|  | Mean SPS | Mean TCS | Notes |
| --- | --- | --- | --- |
| Historian v1.0.2 | 0.822 | 0.497 |  |
| (fast mode) | 0.820 | 0.494 |  |
| ClustalW | 0.787 | 0.447 | From drive5.com |
| Prank v.100701 | 0.784 | 0.441 |  |
| ProbCons v1.12 | 0.883 | 0.619 | From drive5.com |
| Muscle v3.8.31 | 0.843 | 0.532 |  |
| Muscle v4.0 | 0.889 | 0.642 | From drive5.com |

The results of this experiment are summarized in Figure 2. Note that even perfect knowledge of the true evolutionary alignment does not guarantee perfect reconstruction of the underlying indel rate parameters. Indel events can overlap and thus be under-counted, leading to a small negative bias in the estimated rate, and the inherent noisiness of the simulation leads to a spread in the distribution of estimated rates from individual alignments. Historian and Prank (which both explicitly aim to provide ancestral reconstructions) are the most accurate methods for rate estimation, with a slight edge over ProbCons and Muscle. Historian’s default mode has a slight edge over its fast mode in rate estimation accuracy.

As well as varying the rate parameters, we also repeated the benchmark on a symmetric 8-taxon binary tree. This simulation yielded similar patterns: Historian and Prank are still comparable (with a slight edge to Prank in the symmetric tree, *versus* Historian in the GP120 tree), followed by ProbCons, followed by Muscle. As with the tree of Figure 1, varying simulation parameters did not appear to change the ranking.

Table 2 summarizes an evaluation of Historian on the BAliBase 3.0 benchmark, compiled using 3D structural alignments. In general, Historian performs better on these structure-derived benchmarks than Prank, which also performs ancestral reconstruction (Löytynoja and Goldman, 2008). Compared to leading protein aligners that do not attempt ancestral reconstruction, Historian outperforms ClustalW (Larkin *et al.,* 2007), but not Muscle (Edgar, 2004) or ProbCons (Do *et al.,* 2005). No significant difference was found between Historian’s default and fast modes.

**Table 2:** Comparison of runtimes of Historian, Prank and Muscle on the BAliBase 3.0 benchmark.

|  | Average run time per alignment |
| --- | --- |
| Historian v1.0.2 | 233s |
| (fast mode) | 55s |
| Prank v.100701 | 520s |
| Muscle v3.8.31 | 1.9s |

As noted in Table 2, some of the results were taken from the Muscle website, drive5.com/bench/ (Edgar, 2010). To make a direct comparison of runtimes, we re-ran the benchmarks for Prank and Muscle on the same CPU as the Historian benchmark (Intel Xeon 3.20GHz). The runtimes are summarized in Table 3: Historian is an order of magnitude slower than Muscle, but an order of magnitude faster than Prank. Re-running Prank and Muscle resulted in a slight improvement for Prank and a slight deterioration for Muscle compared to the drive5.com data, presumably due to versioning issues (the version of Muscle available for download 3.8.31, whereas the data reported are for version 4.0; conversely, a more recent version of Prank is available than the one benchmarked on the Muscle website). Table 2 reflects our results, where available, and the drive5.com results in other cases.

We did not benchmark MCMC approaches such as BaliPhy (Redelings, 2014), StatAlign (Novak *et al.,* 2008) or HandAlign (Westesson *et al.,* 2012b). These are expected to be more accurate, but generally take much longer. MCMC samplers may be usefully supplemented by decision-theoretic approaches to summarize a sampling run (Herman *et al.,* 2015).Other potential ways to improve accuracy include context-dependent gap penalties, as used by Muscle (Edgar, 2004), and explicit modeling of tandem duplications (Szalkowski and Anisimova, 2013). Incorporating additional data such as structural annotations may further improve alignments (Herman *et al.,* 2014).

## 4 Availability

Historian is available at github.com/ihh/indelhistorian under the CC BY 3.0 US license. Precompiled binaries for Linux or OSX are available. The program may also be compiled from the C++11 source on a POSIX system with Clang (v.6.1.0) and several free libaries (libgsl, libz and Boost). Makefiles to reproduce the analyses reported in this paper are included in the repository, and the simulation data are available at github.com/ihh/gp120sim.

## Acknowledgements

Historian includes code from Ivan Vashchaev (gason), Heng Li (kseq), and the StackOverflow community.

